# Mineralized collagen scaffold pore architecture and glycosaminoglycan content biases anti-inflammatory macrophage phenotype

**DOI:** 10.64898/2026.03.10.710810

**Authors:** Vasiliki Kolliopoulos, Hashni E. Vidana Gamage, Maxwell Polanek, Melisande Wong Yan Ling, Angela Lin, Robert Guldberg, Erik Nelson, Kara Spiller, Brendan A.C. Harley

## Abstract

Macrophages play a central role in early immune response after injury that can shape the success or failure of craniomaxillofacial (CMF) bone repair. While mineralized collagen glycosaminoglycan (GAG) scaffolds have been developed to support osteogenesis, here we define how scaffold pore size, pore alignment, and glycosaminoglycan (GAG) composition influence human monocyte-derived macrophage polarization. We establish flow cytometry, secretome, and gene expression benchmarks to assess primary macrophage polarization toward M1 versus M2 phenotypes in response to cytokine cocktails in 2D culture and 3D scaffolds. We then define the kinetics macrophage polarization in response to scaffold pore architecture and composition in the absence of exogenous cytokines. All scaffold variants support an early pro-inflammatory response followed by a shift toward M2-like phenotypes over seven days reflected by increased CD206 expression, secretion of pro-healing factors such as CCL18, and upregulation of M2a- and M2c-associated genes. Anisotropic scaffolds with smaller pores more robustly drove angiogenic and extracellular matrix related gene expression as well as earlier emergence of M2-like phenotypes. Scaffold GAG chemistry provided an additional tuning mechanism, with chondroitin-6-sulfate variants promoting the greatest late-stage M2 surface marker expression, heparin variants accelerating early M2 and pro-angiogenic phenotypes, and chondroitin-4-sulfate variants dampening both M1 and M2 phenotypes at early timepoints. These findings demonstrate that mineralized collagen scaffolds intrinsically guide macrophage polarization toward pro-regenerative states but that scaffold structure and composition can be used to shape the kinetics and intensity of these responses. These insights provide a critical foundation for immuno-instructive biomaterial designs that enhance CMF bone repair.

## 1. Introduction

Craniomaxillofacial (CMF) bone defects can arise from traumatic injuries, post-oncologic resection, or as a result of developmental defects (e.g., cleft palate defects affect 10.2 per 1000 live births in the US [1, 2]) [3, 4]. They are often large and irregular in shape. Current standard of care prioritizes surgical reconstruction via allograft or autograft, with nearly 500,000 bone graft procedures performed annually in the US alone[5]. Autografts and allografts can suffer from insufficient access to bone and donor site morbidity[6] as well as surgical complications (10-40%) [7–10]. Tissue engineering approaches that combine biomaterials with exogenous cells such as mesenchymal stem cells have been of significant interest due to their potential to induce regenerative healing via multi-lineage differentiation. As a result, a primary biomaterial design criterion has been providing an environment to promote osteogenic differentiation of either exogenously added or endogenously recruited osteoprogenitors [11–13]. However, osteogenic activities must also occur in the context of the robust immune and inflammatory response that exists after injury and biomaterial implantation. Here, hematoma formation after injury triggers the complement system, recruiting monocytes from the blood circulation that differentiate into macrophages and further polarize towards a pro-inflammatory (e.g., M1) phenotype [14, 15].

There is a significant opportunity to pursue biomaterial designs that provide immunomodulatory phenotypes that do not indiscriminately inhibit inflammatory responses but rather help shape a nuanced pro-regenerative environment within the biomaterial implant [16–19]. Notably, pre-inflammatory macrophages contribute to phagocytosis of cellular debris and release cytokines to recruit additional cell populations. For regenerative healing, macrophages must also begin polarizing towards an anti-inflammatory (e.g., M2) phenotype [15]. M2 macrophages work to resolve inflammation by secreting a plethora of anti-inflammatory factors that promote angiogenesis and recruit osteoprogenitors that subsequently differentiate down the osteoblast lineage. The temporal coordination of this sequence is essential for healing. In cases of chronic inflammation, M1 macrophages may not polarize towards an M2 phenotype, leaving a wound environment in a heightened inflammatory stance that halts or inhibits repair [20]. Interestingly, in situations where the initial M1 phenotype is avoided and an M2 response is initiated, tissue repair can also be compromised [21]. As a result, it is paramount to develop strategies to modulate both the magnitude and temporal nature of M1-like to M2-like macrophage transition to support regenerative healing. There is a significant opportunity to understand how biomaterial design may be used to inform macrophage phenotypic progression that aids regenerative healing.

Prior efforts, largely using two-dimensional substrates, have begun to identify biophysical parameters that influence macrophage phenotype. Macrophages seeded on micropatterned surfaces embossed with aligned channels elongate in response to structural anisotropy (alignment) and showed enhanced M2 polarization [22]. Recent literature has begun to extend these observations into 3D, with work showing that 40 μm pores in electrospun meshes can guide macrophage polarization towards an M2 phenotype [23] and that the inclusion of chondroitin sulfate glycosaminoglycans reduces pro-inflammatory cytokine expression in murine macrophages [24]. Biomaterial systems that enable systematic consideration of the effects of microarchitecture and composition would provide important new insight. We are developing a mineralized collagen-glycosaminoglycan scaffold for CMF bone regeneration. This low-density, open-cell foam scaffold was originally optimized to support cell infiltration, MSC osteogenesis and mineral deposition *in vitro*, as well as bone regeneration *in vivo* without the need for exogenously added osteogenic supplements (osteogenic media, BMP-2) [11, 12, 25–35]. However, its porous nature and flexibility in fabrication also provides an avenue to interrogate the role of scaffold architecture and composition on cell signaling in the bone healing microenvironment. We recently reported methods to fabricate variants with control over pore size, pore anisotropy (shape/alignment), and matrix composition. Anisotropic variants containing aligned tracks of ellipsoidal pores promoted cell migration and bone formation [36] while glycosaminoglycan content (chondroitin-6-sulfate, CS6; chondroitin-4-sulfate; heparin sulfate, heparin) was shown to alter MSC osteogenesis and immunomodulatory activity [36–39]. We have begun to examine the activity of macrophage-like cell lines (e.g., THP-1) in these scaffolds [39], motivating efforts here to examine how scaffold structure and composition can influence primary human macrophage polarization in a bone-mimicking microenvironment.

Here, we report the influence of mineralized collagen scaffold pore architecture (aligned vs. non-aligned pores) and glycosaminoglycan content (chondroitin 6-sulfate, chondroitin 4-sulfate, heparin) on the polarization kinetics of primary human macrophages. We culture human primary monocyte-derived M0 macrophages within mineralized collagen scaffolds, quantifying scaffold-induced changes in M1 versus M2 phenotype over a seven-day culture period via shifts in surface marker expression, secreted factors, and gene expression. We establish polarization trajectories for these macrophages using benchmarking data gathered from M1 and M2 macrophages in scaffolds that were polarized with defined cocktails of exogenous factors. The study provides important insight regarding the ability for scaffold architecture and composition to direct the quality and kinetics of M2 polarization while also supporting an initial M1 phenotype essential for regenerative healing.

## 2. Materials and Methods

### 2.1. Experimental design

The study first quantified the effect of known polarizing agents on primary macrophage phenotype in conventional 2D culture and within 3D mineralized collagen scaffolds via established surface marker, secreted factor, and gene expression metrics. Primary human peripheral blood monocytes were first differentiated to M0, M1 (via exogenous LPS+IFNγ), or M2 (via exogenous IL4) phenotypes in conventional 2D culture to benchmark our assessment metrics. For 3D collagen scaffold culture, we also compared already polarized (M0, M1, or M2) macrophages seeded into 3D mineralized collagen scaffolds versus scaffolds seeded with M0 macrophages then exposed to polarizing agents for 2 days. This allowed us to evaluate whether the 3D mineralized collagen scaffold would alter patterns of cytokine driven polarization.

The study subsequently examined the effect of mineralized collagen scaffold glycosaminoglycan composition, pore size and pore alignment on M0 macrophage phenotypic polarization without the use of exogenous polarizing agents. Here, primary human monocytes were polarized to a neutral M0 phenotype, seeded onto each scaffold variant, and then cultured for 7 days in the absence of polarizing factors. Macrophage polarization was assessed via surface marker, secreted factor, and gene expression profiles over the course of a 7-day period, comparing results to the benchmarks for macrophages polarized in the presence of exogenous cytokines.

### 2.2 Fabrication of mineralized collagen-glycosaminoglycan scaffolds

The fabrication of mineralized collagen-glycosaminoglycan scaffolds has been previously described [13, 37, 38]. Briefly, a mineralized collagen precursor suspension containing type I bovine collagen (1.9 w/v% Collagen Matrix Inc., New Jersey USA), calcium salts (calcium hydroxide and calcium nitrate tetrahydrate, Sigma-Aldrich), and one of three potential glycosaminoglycans (0.84 w/v%) – chondroitin 6-sulfate (C6S) sodium salt (CAS Number 9082-07-9, Spectrum Chemicals); chondroitin 4-sulfate (CS4); heparin (Hep) sodium salt from porcine intestinal mucosa (CAS 9041-08-1, Sigma-Aldrich) – were homogenized in a mineral buffer solution (0.1456 M phosphoric acid/0.037 M calcium hydroxide). The precursor solution was then transferred into aluminum molds (isotropic variants) or Teflon molds with a copper plate (anisotropic variants) and lyophilized into porous scaffolds using a Genesis freeze-dryer (VirTis, Gardener, New York USA) [36]. Isotropic scaffolds were generated by cooling the suspensions at a constant rate of 1 ⁰C/min from 20 ⁰C to −10 ⁰C or from 20 ⁰C to −60 ⁰C, followed by a hold at −10 ⁰C or −60 ⁰C for 2 hours, forming larger and smaller ice crystals, respectively [13]. Alternatively, the anisotropic scaffolds were created by freezing the suspension at −10 ⁰C or −60 ⁰C for 2 hours [36]. Following freezing, the frozen suspension was then sublimated at 0 ⁰C and 0.2 Torr, resulting in a porous scaffold interconnected network [40, 41]. A 6 mm diameter biopsy punch (Integra LifeSciences, New Jersey, USA) was used to punch individual isotropic scaffold specimens (6mm dia x 1.5 mm in height), while a razor was used to cut anisotropic 6 mm diameter rods to 1.5 mm in length (6mm dia x 1.5 mm in height).

### 2.3. Characterization of mineralized collagen scaffold pore structure via μCT

Microcomputed tomography (µCT) imaging was performed with a Zeiss Xradia 620 Versa (Carl Zeiss Microscopy, White Plains, New York USA). Individual scaffolds (∼5mm diameter x 3-5mm high) were stacked in syringe tubes (one tube per group). Vertical stitch scanning was used to capture at least 3 full scaffolds for each tube/group, and scan parameters were as follows: 100kV, 15W, exposure 7.3s, filter = Air, Objective = 4X, bin = 1, source position −143.17mm, detector position 13.70mm, 1601 projections, resulting in ∼4.5hr scan time. This produced an isotropic voxel size of 3.09µm. For image analyses, Dragonfly v2021 for Windows (Object Research Systems (ORS), Montréal, Canada, software available at http://www.theobjects.com/dragonfly) with Bone Analysis toolkit was utilized to produce solid and pore microarchitectural quantifications and volume thickness mappings. The automated methodology applied in the Bone Analysis toolkit relies on a dual thresholding technique originally developed for cortical and trabecular bone analyses [42], but as applied to these scaffolds (with no cortex) first an artificial filled cylinder was created as a utility shape surrounding each scaffold and assigned to the ‘cortical’ object, and a ‘filled’ cylinder was created and assigned to the ‘filled’ object. Each scaffold’s associated objects were cropped individually to reduce overall object size and memory required. Solid vs pore (air) were separated via intensity thresholding and kept consistent across scaffolds, and depending on whether the analysis was for solid or pore morphometry, this segmentation (solid or pore) was assigned to the ‘trabecular’ object. An estimated trabecular thickness metric was also required for the Bone Analysis toolkit, and for each individual scaffold analyzed, this was based on preliminary volume thickness and separation data generated for that group. Microarchitectural parameters were calculated based on direct distance transformation methods and included mean intercept length (MIL) anisotropy, star volume distribution (SVD) anisotropy, object volume (mm3), total volume (mm3) object volume fraction (ratio), average strut or pore thickness (µm), and average strut or pore separation (µm) [43–45]. After the computations were completed (∼25-50mins run time), a final volume thickness mapping was created for visual display.

### 2.4. Sterilization, hydration, and scaffold crosslinking

After fabrication all scaffolds were places in sterilization pouches and sterilized via ethylene oxide treatment for 12 hours using a AN74i Anprolene gas sterilizer (Andersen Sterilizers Inc., Haw River, North Carolina USA). All subsequent steps were conducted in aseptic environments. Sterile scaffolds were hydrated and crosslinked using EDC-NHS chemistry as previously described[40, 46–48]. Scaffolds were soaked in 100% sterile filtered ethanol, washed and soaked in PBS, and crosslinked with EDC-NHS. The scaffolds were further washed with PBS and soaked in basal media for 48 hours prior to cell seeding.

### 2.5 Cell culture

#### 2.5.1 Primary human monocyte culture and polarization to classically (M1) and alternatively (M2) activated phenotypes

Primary human peripheral blood monocytes (70034, STEMCELL Technologies, Vancouver, Canada) were cultured, differentiated, and activated into classically (M1) and alternatively activated (M2) macrophages as per the manufacturer’s instructions. Briefly, primary monocytes were plated at a concentration of 1×10^6^ cells/mL into T25 and T175 flasks and were exposed to ImmunoCult™-SF Macrophage Medium (10961, STEMCELL Technologies, Vancouver, Canada) supplemented with 50 ng/mL recombinant human M-CSF (78057, STEMCELL Technologies, Vancouver, Canada). After a 4-day differentiation period, half of the original medium containing M-CSF was added, and the cells were cultured for an additional 2 days (activation period). To generate non-activated (M0) macrophages, no additional cytokines were added to the culture media after the differentiation period. To generate pro-inflammatory/classically activated (M1) macrophages, 10 ng/mL LPS and 50 ng/mL IFNγ were added to the culture media for the 2-day activation period. Anti-inflammatory/alternatively activated (M2) macrophages were generated by adding 10 ng/mL IL4 to the medium for the 2-day activation period.

#### 2.5.2 Lifting macrophages to prepare for seeding

Following the differentiation/activation, excess media was removed from each flask, then centrifuged in 50 ml conical tubes at 300 g for 10 min (a supernatant sample was also retained for ELISA, while the pellet was retained for later use). 5 ml or 10 ml of Accutase was added to each (T25, T175) flask then incubated at 37°C for 15 min. 10 ml or 20 ml of RPMI 1640 supplemented with 10% heat-inactivated FBS and 1% penicillin-streptomycin (complete RPMI media) was used (T25, T175 flasks) to neutralize the Accutase. Cells were then transferred to 50 ml conical tubes and centrifuged at 300 g for 10 min. The supernatant was removed, and cells were resuspended in fresh media at a concentration of 10^6^ cells/ml.

#### 2.5.3 Seeding and culture of human monocyte-derived macrophages on 2D plates and 3D mineralized collagen scaffolds

Macrophages were seeded into 24-well plates for conventional 2D culture. M0, M1, and M2 macrophages were seeded directly on the plate surface at 200,000 cells by placing 20 μl of the stock cell suspension. M1 and M2 groups were subsequently cultured with basal complete RPMI media. M0 macrophage groups were either cultured in basal complete RPMI media (M0 group), supplemented with M1 polarizing cytokines (M0+1), or supplemented with M2 polarizing cytokines (M0+2). All 2D groups were cultured for 48 hours prior to sample collection.

Macrophages seeded on mineralized collagen scaffolds (3D) were cultured in ultra-low attachment 24-well plates, then seeded on each scaffold type at 200,000 cells/scaffold by pipetting 20 μl of the stock cell suspension directly onto the scaffold per an established static-seeding approach [29, 49]. All scaffold groups (Iso, Ani-10, Ani-60, CS4, Hep) were seeded with M0 macrophages and cultured in basal complete RPMI media and cultured for 7 days. A subset of isotropic scaffolds seeded with M0 macrophages were then cultured in either basal complete RPMI media (Iso-M0 group), or supplemented with M1 (M0+1) or M2 polarizing cytokines (M0+2) for 48 hours. Samples were collected on days 1, 3, and 7 for downstream analysis.

### 2.6 Macrophage phenotypic characterization via flow cytometry

#### 2.6.1 Lifting cells from 2D tissue culture surfaces

For all 2D (tissue culture surface) samples, media was removed and samples were washed with 600 μl of PBS. PBS was then removed and samples were exposed to 200 μl TrypLE Express (12605010, ThermoFisher) and incubated at 37 °C for 5 min. TrypLE was neutralized with 400 μl media, and cells were gently scraped using a cell scraper. Each sample was then placed on top of a filter cap in a flow tube to separate cells from any particulates. Samples were centrifuged at 1,100 rpm for 5 min to pellet cells. The supernatant was then removed and cells were resuspended in FACS buffer (00-4222-26, ThermoFisher).

#### 2.5.2 Lifting cells from 3D scaffolds

For all 3D (scaffold) samples, media was removed and samples were washed in 600 μl of PBS. PBS was then removed, and scaffolds were cut into quarters. Scaffolds were then exposed to 600 μl of TrypLE Express and incubated at 37 °C for 5 min on a shaker. To each sample, 600 μl of 5 mg/ml collagenase in HBSS was added at 37 °C for 20 min on a shaker. Next, 1.8 ml of complete RPMI media was added to each sample and collected in a 15 ml conical tube. Each well was further washed with 1 ml of media to collect any remaining cells. The contents of the 15 ml conical tube were added on top of a filter cap in a flow tube to separate cells from any particulates in batches. Samples were centrifuged at 1,100 rpm for 5 min to pellet cells. The supernatant was then removed and cells were resuspended in FACS buffer.

### 2.7 Analysis of macrophage phenotypic markers

Cells were stained with a fixable live/dead marker according to the manufacturer’s instructions (Invitrogen) in PBS for 20 min at 4 °C in the dark. Then cells were washed (5 min, 500 x g at 4 °C) with FACS buffer (PBS containing 2% Fetal Bovine Serum and 1% Penicillin-Streptomycin). Cells were incubated with human Fc blocker in FACS buffer for 10 min at 4 °C and subsequently incubated with flow cytometry antibody specific for human CD80 (B7-1) and CD206 (19.2) in FACS buffer for 20 min at 4 °C in the dark. Antibodies were used at a working dilution of 1:100. The antibody panel is listed in **Supp. Table 1**. After incubation, cells were washed thrice with FACS buffer and fixed with 4% formalin. Flow cytometry was performed on BD LSRII (BD Biosciences) and BD FACSymphony A1 (BD Biosciences). The data were analyzed by FCS Express software. Representative gating strategy is shown in **Supp. Figure 1**.

### 2.8 DNA quantification to determine cell number

DNA was isolated using a DNeasy Blood & Tissue Kit (Qiagen, Hilden, Germany) following the manufacturer’s protocol. Briefly, cell-seeded scaffolds were cut into quarters and placed in 180 mL buffer ATL and 20 μL proteinase K and incubated at 56 °C for 18-24 hours until fully dissolved. Next, 200 μL buffer AL and 200 μL ethanol were added to the lysates and vortexed briefly. The following steps comprised a series of washes and were followed as per the manufacturer’s instructions. DNA concentration was quantified using a NanoDrop Lite Spectrophotometer (Thermo Fisher Scientific) and cell number was determined from a standard curve generated with known cell numbers.

### 2.9 Quantification of secreted factors from macrophages cultured in 2D and 3D

A custom Luminex panel (R&D Systems, Minneapolis, MN) was used to quantify the amount of 13 proteins known to be secreted by M1 and M2 macrophages **Supp. Table 2**. The assay was conducted as per the manufacturer’s protocol, and absorbances were read on a Luminex 200 XMAP system (Luminex Corporation, Austin, TX), and each analyte was normalized to a background control and converted to a concentration using its own standard curve. Expression levels of the analytes are depicted as cumulative concentrations (**Figure 4)**.

### 2.10 RNA isolation and NanoString gene expression analysis

RNA was isolated from cells at Day 0 of the experiment using the RNAqueous™ Total RNA Isolation Kit (Invitrogen, Waltham, MA), then eluted in 30 μl of Elution Solution per established methods [39, 50]. Briefly, RNA was isolated from cell-seeded scaffolds cut into quarters using 1 ml TRIzol (Thermo Fisher Scientific, Waltham, MA) and placed into Phasemaker™ Tubes (Invitrogen, Waltham, MA). Scaffolds with TRIzol were vortexed and incubated at room temperature for 5 minutes [36, 39, 50]. Then, 200 μl chloroform was added to each tube, and scaffolds with TRIzol and chloroform were vortexed and incubated at room temperature for 3 minutes. Tubes were vortexed immediately before centrifuging at 15,000 g for 15 minutes at 4 °C. After centrifugation, the aqueous phase was added to 650 μl 70% ethanol and mixed briefly. This solution was then pipetted into the RNeasy Mini Kit (Qiagen, Hilden, Germany) extraction columns and the kit instructions were followed as per manufacturer’s recommendation. RNA was eluted in RNAse free water and quantified using a NanoDrop Lite Spectrophotometer (Thermo Fisher Scientific, Waltham, MA). RNA samples were stored at −80°C until further analysis.

A custom NanoString panel of 72 mRNA probes was used to quantify transcript expression using the NanoString nCounter System (NanoString Technologies, Inc.; **Supp. Table 3**) [39]. Qubit RNA BR Assay Kit was used to further confirm RNA concentration and quality following thawing. RNA was then loaded into cartridges, and run on the NanoString assay as per the manufacturer’s instructions. The nSolver Analysis Software (NanoString Technologies, Inc.) was used for data processing, normalization, and evaluation of expression. Data is expressed as fold change compared to Day 0 controls on a log2 scale (n=3).

### 2.11 Statistics

Statistical analysis was performed with RStudio (RStudio, Massachusetts, USA) and OriginPro (OriginPro, Massachusetts, USA) software packages. A p-value less than 0.05 was considered statistically significant. Statistics were executed per time point for the scaffold property experiments, and per culture condition (2D vs 3D) for the 2D vs 3D study. The following analysis was performed for Nanostring (n=3), ELISA (n=3), and flow cytometry data (n=5 for experimental groups, n=6 for day 0 controls). First, the assumption of normality and the assumption of equal variances were evaluated using the Shapiro-Wilk normality test and Levene’s test, respectively. If the dataset was both normal and had equal variances, the following was done to assess significance: (i) For the scaffold property experiments and within culture types (2D or 3D) in the 2D vs 3D study, a one-way ANOVA with post-hoc analysis was performed. If data was not normal but had equal variances, a Kruskal-Wallis rank-sum test was performed to assess significance. (ii) For the 2D vs 3D study, a two-way ANOVA with Tukey post-hoc was performed with respect to macrophage state (M0, M0+1, or M0+2) and culture type (2D vs 3D) for the groups replicated in both culture types. However, if data was not normal or had unequal variances, no significance was assessed. All plots were created in OriginPro. Error bars for each group were represented as mean ± standard deviation.

## 3. Results

### 3.1. Benchmarking macrophage polarization in 2D via flow cytometry

Primary monocytes were expanded and differentiated to their non-activated macrophage form (M0) then activated with polarizing cytokines into an M1 or M2 phenotype. We evaluated the effect of polarizing cytokines in 2D culture directly via surface marker expression (M1: CD80+; M2: CD206+; **Figure 1A,B**). M1 polarized macrophages expressed 78% CD80+ and 14% CD206+ while M2 macrophages expressed 0.9% CD80+ and 85% CD206+ (**Figure 1B**). CD206+ expression in M2 polarized macrophages was significantly (p < 0.05) greater than that seen for both non-polarized M0 or polarized M1 phenotypes.

**Figure 1:**
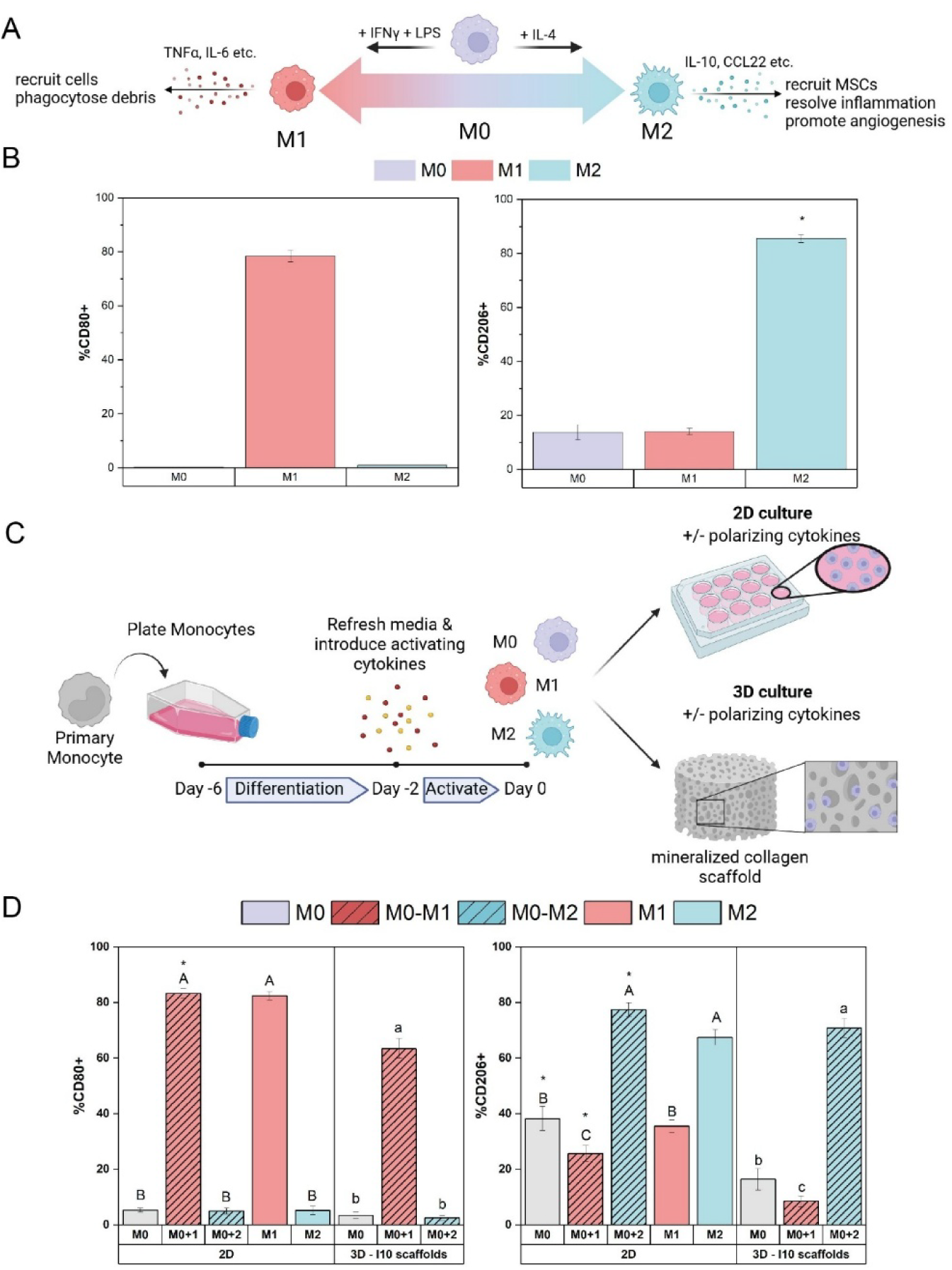
**(A)** Macrophages can take on a gradient of different phenotypes and can largely be described as either non-activated (M0), classically activated/pro-inflammatory (M1) or alternatively activated/anti-inflammatory (M2). **(B)** Pre-polarized macrophages were phenotyped via flow cytometry using CD80+ as the M1 surface marker and CD206+ as the M2 surface marker. * indicated significance of indicated group with all other groups. **(C)** Macrophages were then seeded on 2D plates or 3D scaffolds and exposed to basal media or media supplemented with M1 or M2 cytokines. **(D)** These macrophages were phenotyped via flow cytometry on day 2 of culture. Groups with different capital letters are significantly different in the 2D groups and groups with lower case letters are significantly different in 3D groups. * indicates significance between the 2D and 3D culture methods of the same group.

To evaluate whether the timing of cytokine exposure influences phenotypic expression, unpolarized M0 macrophages were seeded on 2D plates and then cultured in either basal media or exposed to M1 or M2 cytokines (denoted *2D M0+1* or *2D M0+2*, respectively) for two additional days. Alternatively, pre-polarized M1 and M2 macrophages seeded on 2D plates were cultured in basal media for two days (**Figure 1C**). Both M0+1 and M1 macrophages expressed significantly (p < 0.05) greater levels of CD80+ compared to the M0, M0+2, and M2 groups (**Figure 1D**). Both M0+2 and M2 groups displayed significantly (p < 0.05) greater expression of CD206+ compared to the M0, M0+1, and M1 (67-77%) (**Figure 1D**). Interestingly, after two days of culture in basal media both M0 and M1 groups begin to display increased CD206+ surface marker expression. These data suggest polarized macrophages retain similar antigen expression level in the case of immediate polarization after differentiation versus if they are polarized after being seeded in 2D culture as M0 macrophages, confirming the rigor of these benchmarks for assessing macrophage phenotype in scaffolds.

### 3.2. Defining patterns of gene expression and biomolecule secretion for macrophages polarized in 2D culture

We subsequently examined how M1 vs. M2 polarization (by exogenous cytokines) affects short-term changes in macrophage secretome and gene expression in 2D culture. We quantified biomolecule secretion over 7 days, finding exposure of M0 macrophages to M1 polarizing cytokines (2D M0+1) resulted in higher secretion of M1 biomolecules, Tumor Necrosis Factor α (TNFα) and Interleukin-1β (IL-1β), compared to the M0+2 group, although this was not significant (p < 0.05). Looking at specific days, M1 macrophages at day 0 had significantly (p < 0.05) higher secretion of C-C motif Chemokine Ligand 18 (CCL18) than M2 (**Figure 2A-C**). This was reversed on day 2, in which the M1 macrophages exhibited significantly (p < 0.05) lower secretion of CCL18 than the M2 macrophages. After 2 days of culture, M0 macrophages exposed to M1 cytokines (2D M0+1) had similar CCL18 expression as M0 macrophages exposed to M2 cytokines (2D M0+2).

**Figure 2:**
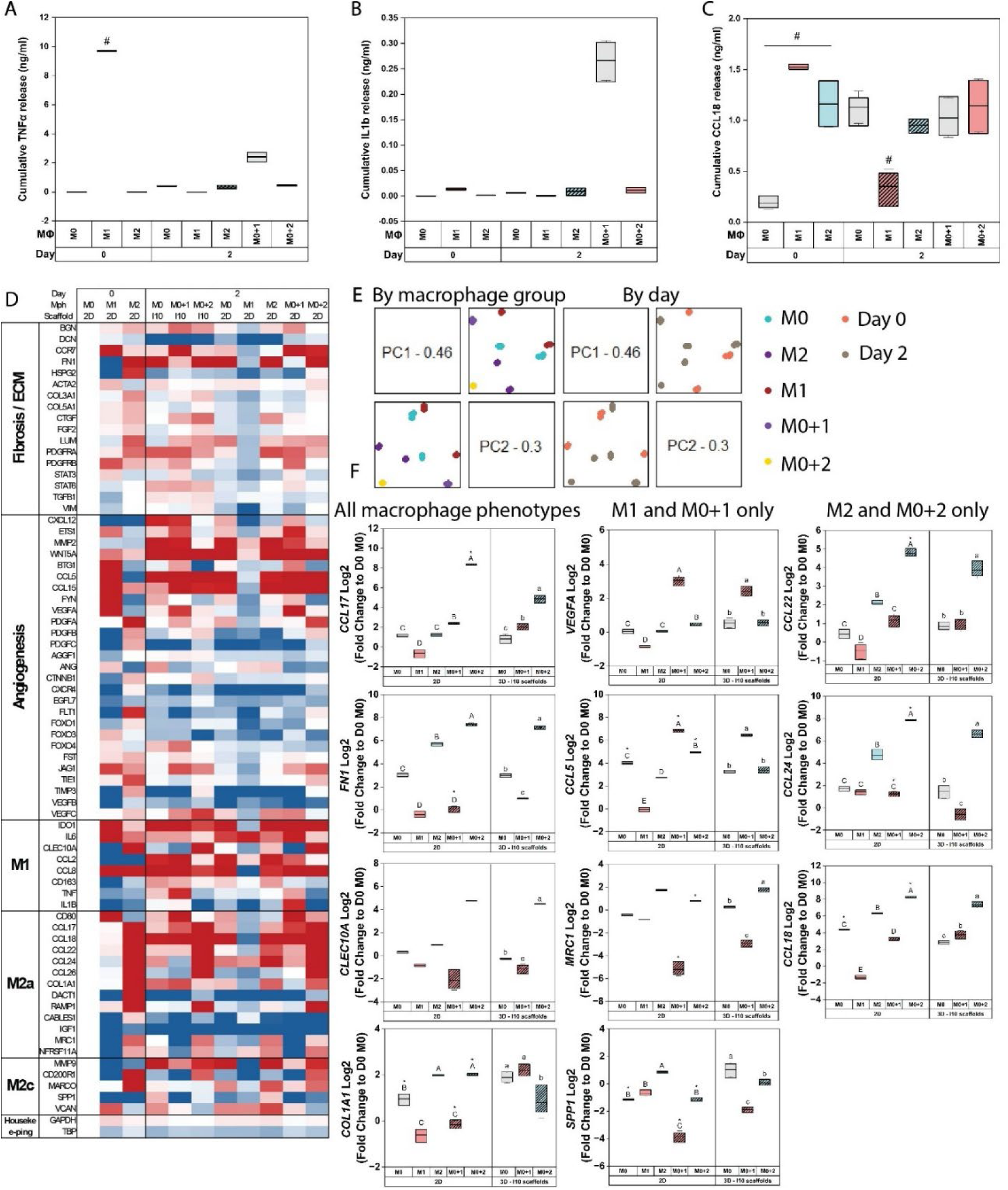
Benchmarking macrophage phenotypic expression in 2D vs in 3D mineralized collagen scaffolds as a function of polarizing cytokines. **(A-C)** TNF-α, IL-1β, and CCL18 ELISA quantification of secreted factors from 2D culture. Day 0 values imply media taken from cell culture in the expansion and differentiation flasks after the activation period. Day 2 values imply seeded macrophages on 2D surfaces either pre-polarized from culture (M1 or M2) or polarized after seeding for 2 days with M1 (M0+1) or M2 (M0+2 cytokines). **(D)** Custom gene expression panel for macrophage relating genes showing fold change expression compared to the M0 day 0 control. Red indicates upregulation while blue indicates downregulation compared to the M0 day 0 control. **(E)** PCA analysis of gene expression as a function of macrophage group and day. **(F)** Differential expression analysis resulted in the identification of 4 genes significantly changing as a function of polarization cytokines shared across all macrophage groups, 4 genes unique to M1 and M0+1 groups, and 3 genes unique to M2 and M0+2 groups.

PCA analysis of gene expression profiles identified shifts in macrophage polarization state as a function of polarizing agents and exposure time (**Figure 2E**). We identified four genes (*CCL17*, *FN1*, *CLEC10A*, and *COL1A1*) that robustly identified M1 versus M2 cytokine-driven polarization (**Tables 1 and 2**; **Figure 2F**), with all four upregulated (relative to M0) in M2 and M0+2 conditions, and all but CCL17 downregulated (relative to M0) for M1 and M0+1 polarization. Interestingly, while CCL17 was downregulated in M1 pre-polarized cells it was upregulated in M0+1 cells. Further, pre-polarized M1 macrophages and M0 macrophages exposed to M1 cytokines (2D M0+1) share 12 genes that are significantly different (p < 0.05) versus M0 macrophages, with *VEGFA* and *CCL5* upregulated and all others downregulated. Pre-polarized M2 macrophages and M0 macrophages exposed to M2 cytokines (2D M0+2) shared seven common significantly upregulated genes (vs. M0 control), with *CCL24*, *CCL17*, *CCL22*, *CCL18*, and *CLEC10A* also more significantly upregulated in the M0+2 group. Overall, this identified a rigorous set of gene and surface antigen expression metrics to differentiation M0, M1, and M2 polarization state for subsequent scaffold studies.

**Table 1.** Gene expression shifts in response to M1 polarizing cytokines. Fold change at day 7 of culture reported as relative to M0 macrophages.

|  | **2D M1**<br>**Fold Change** | **2D M1**<br>**p-value** | **2D M0+1**<br>**Fold Change** | **2D M0+1**<br>**p-value** |
| --- | --- | --- | --- | --- |
| *VEGFA* | 2.20 | 0.000639 | 3.46 | 9.02e-05 |
| *CCL5* | 2.70 | 0.00352 | 3.38 | 0.00441 |
| *CCL17* | -2.73 | 0.000767 | 2.29 | 0.000915 |
| *SPP1* | -3.06 | 0.000114 | -5.47 | 2.41e-06 |
| *MRC1* | -3.94 | 0.000399 | -6.76 | 1.81e-05 |
| *FN1* | -3.86 | 2.48e-06 | -3.31 | 0.000136 |
| *CD200R1* | -3.21 | 0.000185 | -4.27 | 0.000165 |
| *CLEC10A* | -2.27 | 0.00263 | -3.02 | 0.00216 |
| *COL1A1* | -2.96 | 0.000394 | -3.32 | 0.00218 |
| *HSPG2* | -2.64 | 0.000193 | -2.95 | 0.00745 |
| *DACT1* | -2.01 | 0.0078 | -2.14 | 0.0316 |
| *PDGFA* | -3.13 | 0.00117 | -2.24 | 0.0396 |

**Table 2:** Gene expression shifts in response to M2 polarizing cytokines. Fold change at day 7 of culture reported as relative to M0 macrophages.

| *com* | **2D M2**<br>**Fold Change** | **2D M2**<br>**p-value** | **2D M0+2**<br>**Fold Change** | **2D M0+2**<br>**p-value** |
| --- | --- | --- | --- | --- |
| CCL22 | 6.34 | 1.73e-08 | 7.18 | 1.13E-07 |
| FN1 | 5.64 | 1.79e-09 | 5.03 | 4.61E-07 |
| CCL24 | 4.05 | 4.25e-13 | 6.67 | 3.79E-16 |
| CCL18 | 4.37 | 1.93e-06 | 4.83 | 1.41E-05 |
| CCL17 | 2.09 | 0.000238 | 8.66 | 9.2E-12 |
| CLEC10A | 2.8 | 0.000385 | 4.37 | 5.31E-05 |
| COL1A1 | 2.97 | 1.16e-07 | 2.12 | 0.000117 |

### 3.3. Mineralized collagen scaffolds do not pose a diffusion barrier for cytokine-driven polarization

To evaluate whether there are shifts in macrophage phenotype as a result of 3D scaffold culture, M0 macrophages were seeded into scaffolds then cultured in basal media or exposed to M1 or M2 polarizing cytokines after seeding (3D M0+1, 3D M0+2) (**Figure 1C**). In the 3D scaffold groups, the M0+1 group displayed significantly (p < 0.05) greater expression of CD80+ (60%) than M0 or M0+2 groups (**Figure 1D**). While still predominantly expressing CD80, CD80 expression was lower (p < 0.05) in macrophages polarized in 3D scaffolds versus to 2D culture (Figure 1D). Similarly, 3D M0+2 macrophages displayed significantly (p < 0.05) greater CD206+ expression compared to 3D M0 or M0+1 groups (Figure 1D), with different levels in CD206 expression (p < 0.05) between 2D vs 3D culture methods. Macrophages were seeded on scaffolds then stimulated with M1 polarizing cytokines (3D M0+1) showed an increase in IL1b and TNFα secretion, suggesting a more pro-inflammatory state (**Figure 2A,B**). Analysis via a custom Luminex assay identified an increase in IL-6, CXCL1, CXCL2, and CXCL9 secretion in 2D M1, 2D M0+1 and 3D M0+1 groups as well as an increase in CCL17 in 2D M0+2 and 3D M0+2 groups (**Supp. Figure 2**).

Seeding macrophages on scaffolds also affected gene expression (**Figure 2F**). Expression of the M1 gene *IL6* was reduced by 3D culture among M0 and M0+1 macrophages (p < 0.05), while M1-associated *TNF* was significantly increased in M0, M0+1 and M0+2 groups in 3D compared to 2D (p < 0.05). Macrophages in 3D scaffolds exhibited increased angiogenic genes *CTNNB1* and *VEGFB* versus 2D culture for M0, M0+1 and M0+2 groups (p < 0.05). The expression of several M2a genes (*CCL17*, *CCL18*, *CCL24*, and *COL1A1*) was dampened in 3D vs. 2D culture for the M0+2 groups. Interestingly, expression of M2a genes *COL1A1* and *MRC1* was heightened for M0+1 macrophages in 3D scaffold culture, surpassing expression seen for M0+2 macrophages in 3D. Similarly, expression of angiogenic gene *PDGFB* and fibrosis/ECM gene *HSPG2* was higher in M0+1 macrophages versus M0+2 macrophages in 3D, suggesting the importance of initial M1 polarization in long-term pro-healing phenotype. Expression of fibrosis-associated *CCR7* and *DCN* was significantly dampened for both M0+1 and M0+2 when cultured in 3D compared to 2D (p < 0.05). Taken together, these data suggest that while the effect might be slightly blunted, macrophages can be effectively differentiated in 3D scaffolds towards M1 and M2 phenotypes using conventional cytokine cocktails and that scaffold culture may induce additional pro-healing phenotypes..

### 3.4. Mineralized collagen scaffold pore architecture analysis via μCT

We then examined the role of scaffold structure on macrophage phenotype, initially assessing changes in scaffold architecture as a result of lyophilization (fabrication) conditions. μCT analysis and an advanced analysis tool, Dragonfly, was used to evaluate the degree of anisotropy and pore thickness in scaffolds as a result of directional (anisotropic) vs. isotropic heat transfer during lyophilization (**Figure 3**). Anisotropic (A) scaffolds fabricated at higher (−10°C freezing: (A10) and lower (−60°C freezing: A60) have significantly larger degree of anisotropy compared to isotropic (I) scaffolds fabricated at the same freezing temperatures (I10, I60). Isotropic scaffolds fabricated at a higher freezing temperature (−10°C) had the largest pores compared to all groups, including anisotropic scaffolds fabricated at the same freezing temperature.

**Figure 3:**
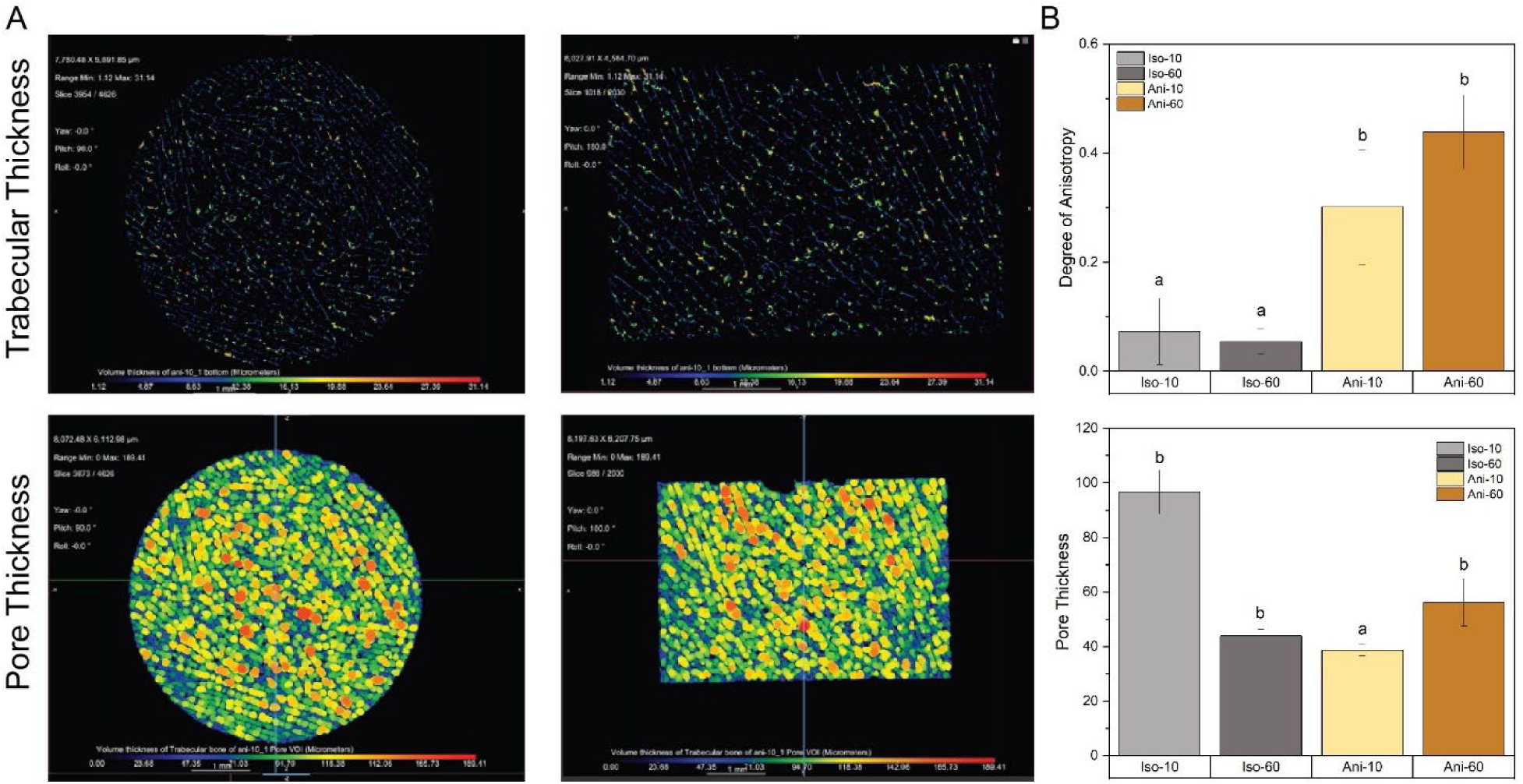
Quantification of scaffold pore architecture via μCT analysis. **(A)** Representative images of the scaffold in Dragonfly software. **(B)** Degree of anisotropy and pore thickness quantified by Dragonfly. Anisotropic scaffolds (A10, A60) have significantly higher pore anisotropy compared to isotropic scaffolds (I10, I60). Isotropic scaffolds at higher freezing temperature (−10 °C) have significantly larger pores compared to anisotropic scaffolds at the same temperature. Groups with different letters indicate significance (p <0.05).

### 3.5. Scaffold pore size enhances M2 macrophage surface marker expression

After defining benchmarks of M1 vs. M2 polarization in response to polarizing cytokines, we then defined the effect of scaffold microarchitecture on macrophage polarization in the absence of exogenous polarizing agents. Three scaffold variants were examined: our conventional variant fabricated with isotropic pores at a freezing temperature of −10°C (I10); an anisotropic (aligned) variant fabricated at a freezing temperature of −10°C (A10); or an anisotropic (aligned) variant fabricated at a lower freezing temperature of −60°C (A60) that exhibited a significantly smaller pore size. M0 macrophages were seeded on scaffolds and cultured for 7 days in the absence of polarizing agents (**Figure 4A**). M0 macrophages seeded on isotropic scaffolds (I10) and anisotropic scaffolds with large or small pores (Ani10 and A60 respectively) displayed low (<5%) expression levels of M1-associated CD80+ throughout all 7 days of culture (**Figure 4B**). By day 7 of culture, macrophages in small pore anisotropic (A60) scaffolds displayed significantly (p < 0.05) lower CD80+ expression than conventional isotropic variants (I10). While on days 1 and 3 of culture, macrophages in all scaffold variants (I10, A10, A60) expressed similar levels of M2-associated CD206+ (20%), by day 7 macrophage cultures in conventional isotropic scaffolds (I10) expressed significantly (p < 0.05) higher CD206+ levels (50% CD206+) than anisotropic (aligned) A60 variants (28% CD206+) (**Figure 4C**). Taken together, this indicates that larger pore scaffolds (I10 and A10) induce greater amounts of M2-associated surface marker expression in macrophages with a minimal effect of anisotropy.

**Figure 4:**
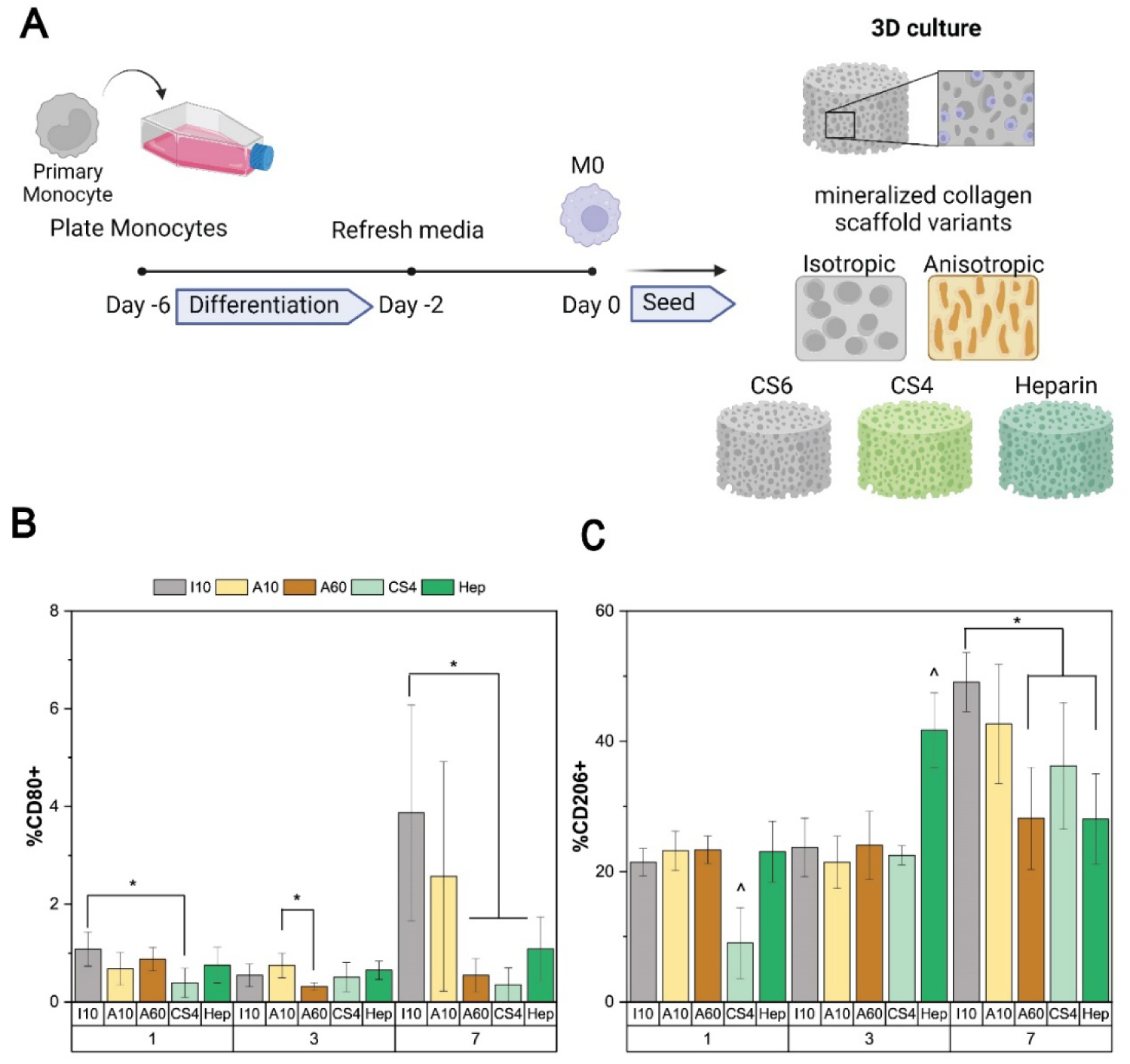
**(A)** Monocytes differentiated to M0 macrophages over 6 days of culture and seeded on mineralized collagen scaffolds with varying pore size and anisotropy or varying glycosaminoglycan type for 7 days of culture. **(B)** M1 macrophage surface marker, CD80, and **(C)** M2 macrophage surface marker, CD206, quantified via flow cytometry (n=3) for each scaffold condition. * indicates significance at p < 0.05 between indicated groups at that timepoint, while ^ indicates significance at p < 0.05 between all other groups at that timepoint.

### 3.6. Scaffold pore architecture does not significantly influence soluble factor expression

We subsequently examined secretion of CCL18, IL1b, and TNF-α as a function of scaffold architecture (**Figure 5**). While no significant differences were observed as a function of structure, the A60 scaffold group displayed increased IL1b expression, suggesting a heightened inflammatory state. Interestingly, seeding M0 macrophages on the mineralized collagen scaffolds induced increased CCL18 factor release with time, suggesting the scaffolds display a potential for temporal polarization.

**Figure 5:**
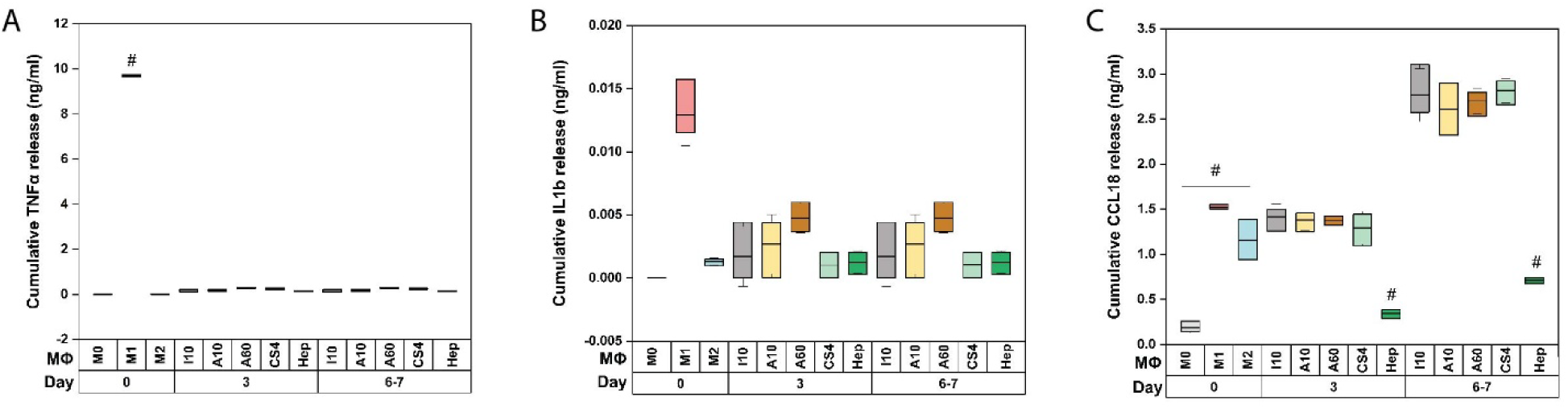
Quantification of secreted factors released by M0 macrophages seeded on mineralized collagen scaffolds as a function of scaffold structure and composition. Cumulative release profiles of (A) TNF-α, (B) IL-1β, and (C) CCL18. # indicates significant differences between

### 3.7. Scaffold pore size enhances M1 macrophage gene expression

Broadly, seeding macrophages on scaffolds drives an initial pro-inflammatory response followed by a progression toward an M2-like gene expression state (**Figure 6**). Cells cultured on I10, A10, and A60 scaffolds all exhibited significant upregulation of 6 genes compared to M0 macrophage at day 0 (**Table 3**), including two genes (SPP1 and CCL5) associated with polarized M1 and M0+1 macrophages from our benchmarking 2D study. None of the altered genes in the 3D scaffold groups overlapped with the notable M2 and M0+2 genes from the 2D study. However, PCA analysis revealed macrophage gene expression profiles within mineralized collagen scaffolds exhibit a temporal shift from an initial pro-inflammatory state to a transition away from that M1 state.

**Figure 6:**
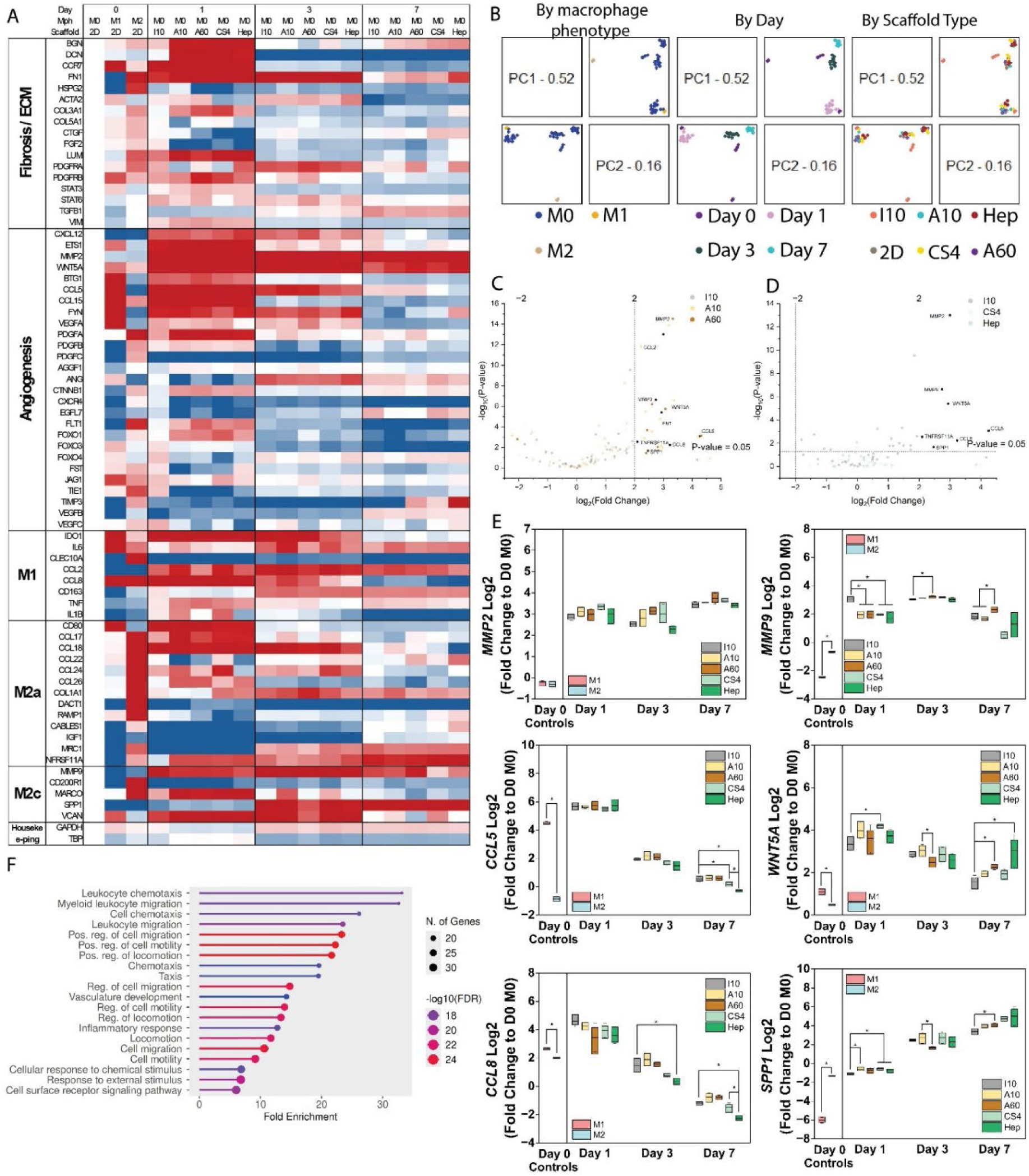
Gene expression evaluation via a custom Nanostring panel. **(A)** Heatmap of macrophage-related genes showing fold change expression compared to the M0 day 0 control. Red indicates upregulation while blue indicates downregulation compared to the M0 day 0 control. **(B)** PCA analysis of gene expression as a function of macrophage phenotype, scaffold type, and day. Volcano plots of −log10(p-value) plotted against log2(fold change) for each gene as a function of **(C)** structure and **(D)** composition scaffold group. Thresholds for both p-value and fold change were set a 0.05 and 2 respectively. Genes above these thresholds are bolded while below the thresholds are transparent. **(E)** Highly upregulated genes in scaffolds groups compared to M0 day 0. **(F)** GO ontology enrichment analysis using The ShinyGO Pathway Analysis to illustrate the top 20 pathways in the comparison performed. All differentially expressed genes (DEGs) with a significant level (FDR value for DEG less than 0.05) are mapped to the referential canonical pathways available in the Culex quinquefasciatus database. The size of a circle represents the number of DEGs classified into one specific pathway category. Fold Enrichment (FE) indicates the percentage of DEGs belonging to a pathway, divided by the corresponding percentage in the background. FDR for FE demonstrates how likely the enrichment is by chance.

**Table 3:** Genes identified in response to macrophage culture on scaffolds with different pore structures.

|  | *I10 Fold Change* | *I10 p-value* | *A10 Fold Change* | *A10 p-value* | *A60 Fold Change* | *A60 p-value* | *2D M1, M0+1* |
| --- | --- | --- | --- | --- | --- | --- | --- |
| *MMP2* | 3 | 9.72E-14 | 3.19 | 1.29E-14 | 3.33 | 2.96E-15 |  |
| *MMP9* | 2.74 | 2.30E-07 | 2.39 | 3.30E-06 | 2.61 | 6.22E-07 |  |
| *WNT5A* | 2.94 | 4.01E-06 | 3.37 | 2.79E-07 | 3.07 | 1.79E-06 |  |
| *CCL5* | 4.25 | 0.000846 | 4.21 | 0.000935 | 4.31 | 0.00073 | Upregulated |
| *SPP1* | 2.47 | 0.0208 | 2.93 | 0.00671 | 2.81 | 0.00904 | Downregulated |
| *CCL8* | 3.23 | 0.00574 | 2.97 | 0.0107 | 2.41 | 0.0362 |  |

Examining the effect of scaffold architecture, anisotropic (aligned) scaffolds (A10 and A60) drove significant (p < 0.05) increased expression of angiogenic genes *CTNNB1*, *PDGFA*, and *VEGFB* compared to isotropic (I10) scaffolds. Anisotropic scaffolds also induced increased expression of genes associated with fibrosis and ECM deposition (BGN, *DCN*, *FN1*, and *STAT3*). Anisotropy and pore size influenced macrophage polarization dynamics. There was a less notable influence of anisotropy and pore size on M1 polarization; nonetheless, on day 3 of culture, anisotropic scaffolds with larger pores (A10) drove significantly (p < 0.05) higher levels of M1-assoicated *IL6* compared to anisotropic scaffolds with smaller pores (A60) and higher levels of M1-associated *CCL2* compared to both anisotropic scaffolds with smaller pores (A60) as well as isotropic scaffolds with larger pores (I10). However, scaffold anisotropy drove significantly (p < 0.05) higher expression of M2 genes than isotropic (I10) scaffolds on day 1 after seeding, including notable M2a genes *CCL17* (A10 only), *CCL26, IGF1*, *DACT1* (A60 only), and *TNFRSF11A* (both), as well as M2c genes *MARCO* and *SPP1* (A10 only). Similar trends were observed at later culture times for M2-associated genes, with both anisotropic (aligned; A10 and A60) scaffolds driving significantly (p < 0.05) higher expression of M2a-associated *CCL18* (day 3) and *COL1A1* (day 3) than the isotropic (I10) scaffold. Anisotropic (aligned) scaffolds with the smaller pore size (A60) also drove significantly higher expression of M2c genes *MMP9* (days 3 and 7) and *SPP1* (day 7) than isotropic (I10) scaffolds.

### 3.8. Scaffold glycosaminoglycan content can instruct M2-like macrophage polarization

We then examined the effect of scaffold glycosaminoglycan (GAG) content on macrophage polarization in the absence of exogenous polarizing agents. All GAG variants were fabricated as identical isotropic (non-aligned) scaffolds at −10°C, matching the isotropic (non-aligned) I10 variant from the earlier comparison of scaffold pore architecture on macrophage polarization. The I10 variant was also used because it is the long-standing “base” variant that shows exceptional osteogenic activity in prior in vitro and in vivo studies. While this base scaffold is traditionally fabricated using chondroitin-6-sulfate (CS6) we compared macrophage polarization in I10 variants created with chondroitin-4-sulfate (CS4) or heparin (Hep) as the GAG content for up to 7 days of culture (**Figure 4**). While M0 macrophages seeded into all GAG variants displayed low M1-associated CD80+ levels (<5%), those seeded into CS4 scaffolds displayed significantly (p < 0.05) lower levels of CD80+ expression compared to C6S on days 1 and 7 (**Figure 4B**). Interestingly, M0 macrophages seeded into CS4 scaffolds also displayed significantly (p < 0.05) lower levels of M2 associated CD206+ after 1 day of culture. By day 3 of culture, M0 macrophages seeded into Hep scaffolds displayed significantly (p < 0.05) higher levels of M2-associated CD206+ than C6S or C4S variants (41%) (**Figure 4C**). By day 7 of culture, a more M2-like phenotype was observed across the board, with macrophages cultured in C6S scaffolds displaying the highest level of M2-associated CD206+, significantly (p < 0.05) higher than those cultured in Hep scaffolds.

### 3.9. Chondroitin sulfate-containing scaffolds have greater anti-inflammatory macrophage-mediated secretome expression

While few differences in macrophage secretome were observed as a function of scaffold glycosaminoglycan content, conditioned media generated over 7 days by macrophages cultured in heparin (Hep) scaffolds contained significantly reduced expression of M2a-associated CCL18 compared to macrophages cultures in chondroitin-4-sulfate (CS4) or our baseline chondroitin-6-sulfate (CS6, I10) scaffolds (**Figure 5**). Using a custom Luminex assay we identified that macrophages cultured in Hep scaffolds also secreted reduced of inflammatory (M1-assocaited) CXCL2 and anti-inflammatory (M2-associated) CLL2 compared to macrophages in either chondroitin sulfate (CS4, CS6/I10) scaffolds by Day 1 (**Supp. Figure 3**). Interestingly, after 3 and 7 days macro[phages in both CS4 and Hep scaffolds secreted increased levels of inflammatory IL-6. Taken together, these data suggest that the inclusion of CS6 or CS4 in mineralized collagen scaffolds increases the anti-inflammatory potential of macrophages. Given the presence of GAG content within the scaffold, this may be via either changes in macrophage surface receptor activation or sequestration of biomolecules via different levels of GAG-sulfation.

### 3.10. Mineralized collagen scaffold promotes a pro-healing macrophage secretome

Analysis of gene expression profiles for M0 macrophages seeded in scaffolds with disparate GAG content revealed significant upregulation of 6 genes compared to the M0 day 0 control (**Table 4**). Notably, two genes (SPP1 and CCL5) were consistent with those seen in cytokine differentiated M1 and M0+1 genes from the 2D study while none were in common with those M2 and M0+2 groups from the 2D study. Based on PCA analysis, all GAG scaffold variants induce an initial M1-like polarization followed by a transition away from that M1 state (**Figure 6**). Looking at changes in time, chondroitin-sulfate (CS4 or CS6) variants supported an initial M1 gene expression phenotype, driving significantly (p < 0.05) higher expression of M1-associated *CCL2* (day 3, CS4 only), *CLEC10A* (day 7, CS6 only) and *IDO1* (day 3, CS6 only) vs. Heparin variants. Meanwhile, CS4 and Heparin variants induced significantly (p < 0.05) higher expression of M2-associated genes on day 1 compared to CS6 variants, including M2a associated *CCL17*, *CCL26*, *DACT1*, and *TNFRSF11A* as well as M2c associated *MARCO* and *SPP1*.

**Table 4:** Genes identified in response to macrophage culture on scaffolds with different glycosaminoglycans.

|  | *I10 Fold Change* | *I10 p-value* | *CS4 Fold Change* | *CS4 p-value* | *Hep Fold Change* | *Hep p-value* | *2D M1, M0+1* |
| --- | --- | --- | --- | --- | --- | --- | --- |
| *MMP2* | 3 | 9.72E-14 | 3.38 | 1.73E-15 | 2.98 | 1.31E-13 |  |
| *MMP9* | 2.74 | 2.30E-07 | 2.27 | 8.13E-06 | 2.25 | 9.09E-06 |  |
| *WNT5A* | 2.94 | 4.01E-06 | 3.44 | 1.75E-07 | 3.36 | 3.01E-07 |  |
| *CCL5* | 4.25 | 0.000846 | 4.06 | 0.00138 | 4.28 | 0.000791 | Upregulated |
| *SPP1* | 2.47 | 0.0208 | 3.53 | 0.00131 | 3.81 | 0.000574 | Downregulated |
| *CCL8* | 3.23 | 0.00574 | 2.59 | 0.025 | 2.24 | 0.0509 |  |

We then examined a broader set of regenerative phenotypes. M0 macrophages seeded into CS4 and Heparin scaffolds showed rapid angiogenic and fibrotic responses, with these variants driving significantly (p < 0.05) higher expression of angiogenic (*CTNNB1*, *PDGFA*, *VEGFB*) and fibrotic (*BGN*, *DCN*, *FN1* and *STAT3*) genes compared to CS6 variants (**Figure 6**). Heparin scaffolds also drove higher expression of angiogenic genes *PDGFC* and *WNT5A* expression on day 1 compared to CS6 variants. Overall, Heparin variants often had higher regenerative gene expression in the late stage of culture compared to chondroitin-sulfate (CS4 or CS6) variants, including angiogenic genes *PDGFB* (day 3, vs. CS4), *PDGFC* (day 3), *FYN* (day 3, vs. CS4; day 7, vs. CS6), and *WNT5A* (day 7, vs. CS6), as well as M2a genes *CD80* (day 7) and *TNFRSF11A* (day 7, vs. CS6) and M2c gene *CD200R1* (day 3).

### 3.11. Gene Ontology analysis reveals high association with chemotaxis and migration

Lastly, we considered how observed shifts in gene expression patterns may inform a broader pro-regenerative stance, using ShinyGO Pathway Analysis to identify the top 20 pathways associated with significantly different genes in M0 macrophages cultured in mineralized collagen scaffolds (versus the day 0 M0 control; **Figure 6**). This analysis identified highly enriched pathways associated with chemotaxis, migration, and motility and suggests that regardless of scaffold GAG content or pore architecture, exposure to this class of mineralized collagen scaffold instructs M0 macrophage motility and chemotaxis driven migratory behavior, factors essential in early wound healing.

## 4. Discussion

Macrophages are a key immune cell type in the bone healing cascade and can exist on a spectrum of phenotypes ranging from pro-inflammatory (M1) to anti-inflammatory (M2). These cells have complex temporal responses in the wound microenvironment that orchestrate whether the tissue will regenerate or if it will remain in a chronic inflammatory state. As a result, biomaterial design motifs able to instruct induce shifts in macrophage phenotype are likely as essential as those able to instruct mesenchymal stem cell osteogenic differentiation for achieving robust regenerative healing. As a result, tunable, biomimetic scaffold systems offer a path to examine the influence of biomaterial structure and composition on macrophage phenotypic response, particularly the capacity to selectively polarize macrophages towards a pro-regenerative M2 phenotype.

We first demonstrated exposing human peripheral blood mononuclear cell derived M0 macrophages to known M1 and M2 polarization cytokines drove quantifiable shifts in phenotype similar to those seen for pre-polarized M1 and M2 macrophages then seeded into scaffolds. These studies used experimental benchmarks established by 2D macrophage culture experiments to assess macrophage phenotype within the 3D scaffold environment, similar to prior observations from Spiller et. al. that found exposing macrophages to inflammatory cytokine combinations in 2D culture can induce distinct polarization states while also demonstrating the potential for unique macrophage responses following an M1-M2 transition [51, 52]. Spiller et. al. also showed macrophages exposed to polarizing cytokines at different times and to either M0 or M1 macrophages drove distinct gene expression profiles and the emergence of a hybrid M1/M2 phenotype as well as a subsequent shift towards an M2 state. As a result, we first evaluated the baseline response of M0 macrophages to polarizing cytokines in 2D, exposing macrophages to the cytokines either prior to or after seeding onto 2D culture flasks. We observed that exposing M0 macrophages to M1 or M2 polarizing cytokines in 2D culture induced increased expression levels of M1 surface markers (CD80+) or M2 surface markers (CD206+), respectively, compared to the pre-polarized (M0) groups. We then wanted to investigate whether the scaffolds would pose a diffusion barrier for cytokine driven-polarization, but observed the same trend for M0 macrophages seeded into 3D mineralized collagen scaffolds when exposed to polarizing cytokines as we saw in 2D,. Together, this confirms polarizing cytokines are potent directors of macrophage phenotypic transition and that the 3D mineralized collagen scaffold does not natively inhibit the effect of polarizing cytokines on macrophage state.

We then subsequently examined if the mineralized collagen scaffold itself provides instructive signals sufficient to regulate macrophage polarization in the absence of conventional cytokines. Broadly, we found that the mineralized collagen scaffold variants collectively enhance the eventual transition of M0 macrophages towards an anti-inflammatory M2 phenotype by first eliciting a pro-inflammatory M1 response. This aligns with our prior findings investigating the polarization of the THP-1 macrophage cell line in chondroitin-6-sulfate isotropic (C6S/I10) mineralized scaffold variant, which also revealed M2-like polarization with time in the absence of exogenous stimulation [39]. While the chondroitin-6-sulfate isotropic (C6S/I10) mineralized scaffold variant was developed to instruct MSC osteogenic differentiation, prior work has begun to explore the effect if additional design modifications on cell activity. Mineralized collagen scaffolds are fabricated by a lyophilization process where control over freezing conditions influences the size and shape of the ice crystals that form in an aqueous collagen-GAG suspension [53], resulting in changes in the size and shape of the resultant pores after sublimation. Notably, reducing the freezing temperature accelerates freezing and the formation of smaller ice crystals then pores[40, 54] while a directional solidification process creases aligned tracks of anisotropic pores versus isotropic scaffolds [55–58]. Further, changing the GAG content in the collagen-GAG suspension alters scaffold composition and can differentially regulate biomolecule sequestration and bioavailability [38, 59]. While we have reported the effects of these modifications on mesenchymal and adipose derived stem cell differentiation [11, 27, 38, 55], emergence of literature suggesting macrophages are mechanosensitive motivated our effort the define the role of scaffold pore architecture (pore size, alignment) and glycosaminoglycan content (CS6, CS4, Hep) on the kinetics of macrophage polarization. Gene expression analysis and principal component analyses revealed that after day 1 of culture, M0 macrophages seeded on mineralized collagen scaffold groups displayed a more pro-inflammatory, M1-like phenotype, while at later timepoints they transitioned away from that inflammatory stance. Across all scaffold variants we observed an initial increase in pro-inflammatory genes (*CCL5* and *CCL8*) that decreased with longer culture. We also observed an increase in remodeling and pro-healing associated genes (*SPP1*, *MMP2*, and *MMP9*) with culture time. When comparing the effect of pore size and anisotropy, we observed that anisotropy had a greater effect than pore size on macrophage gene expression, with anisotropic variants displaying significantly greater expression of angiogenic, ECM associated, and M2-like genes compared to the base (isotropic) I10 variant. Analysis of macrophage surface marker expression suggested that M0 macrophages seeded into scaffolds with larger pore sizes regardless of shape (I10 or A10 variants) displayed increased M2-like phenotype. It should be noted that regardless of pore isotropy or anisotropy, all scaffold variants have a pore architecture defined by networks of collagen-GAG fibers (struts) that each have a large aspect ratio. Notably, cells in fiber networks with large pore sizes are more likely to elongate along a single fiber as opposed to achieving a spread morphology attached to multiple adjacent fibers [60–62]. As a result, our observation that scaffolds with large pores induced M2-like shifts in macrophage phenotype aligns well with recent literature suggesting 2D substrates that induce macrophage elongation also precipitate an M2-like response [23, 63].

All glycosaminoglycan variants induced M2 like polarization, with the effect after 7 days in culture most prominent in isotropic scaffolds containing chondroitin-6-sulfate (C6S/I10). CD206 (M2-like) expression far outweighed CD80 (M1-like) expression at all timepoints and in all GAG groups, suggesting mineralized collagen scaffolds broadly promote M0 to M2 phenotypic transition over the course of a week in culture. Interestingly, macrophages in heparin containing scaffolds both secreted the least soluble CCL18 (a marker of M2-transition) but also exhibited the greatest reduction of M1-associated genes across the culture period. Others have also shown that chondroitin sulfate has potent anti-inflammatory activity towards macrophages by limiting NF-κB signaling [24, 64] while heparin sulfate’s increased ability for charge-based biomolecule sequestration can create local concentration gradients and accumulation of signaling molecules biasing macrophage polarization [65]. Taken together, our studies suggest that biomolecule sequestration by heparin sulfate containing scaffolds may locally amplify M1-associated signaling, resulting in the greatest dampening of M1-associated genes, but may also reduce the amount of available M2-associated polarizing factors.

Interestingly, in one of the earliest examples of tissue regeneration, the GAG content of a non-mineralized collagen-GAG scaffold was shown to significantly impact the kinetics of wound contraction and quality of wound healing, with CS6-containing scaffolds promoting the most significant regeneration of full-thickness skin wounds [66]. As a result, it is likely important to understand the role of multi-cellular interactions in the wound environment. Our findings here clarifying the instructive nature of collagen scaffold pore architecture and GAG content on human macrophage polarization is critical. We establish benchmarks for profiling macrophage phenotype in bone-mimetic 3D scaffolds via surface antigen, genomic, and secretome markets. These results now provide the basis for future efforts to examine crosstalk between macrophages and hMSCs within the scaffold as well as to examine the influence of exogenous inflammatory signals associated with the craniofacial bone microenvironment. While conventional macrophage cell lines provide batch-to-batch consistency, biomimetic scaffolds provide an environment to examine donor variability in cell phenotype. We recently reported a strategy to assess hMSC and adipose derived stem cell (hASC) donor variability in proliferation and osteogenic capacity in the mineralized collagen scaffold [50]. This revealed significant discrepancies in cell metabolic activity, proliferation, and osteogenic capacity, some of which were associated with donor-reported sex. Integrating these studies now using donor-derived hMSCs and human monocytes are likely to provide critical, actionable data regarding strategies to increase osteogenic-immunomodulatory coupling at the implant-wound margins critical for improved regenerative healing. Indeed, for a portion of our study of macrophage polarization in collagen scaffolds here, we examined differences between two donor-derived monocyte specimens. Given the cost-prohibitive nature of expanding and testing in parallel many donor-derived monocyte lines, we only examine a limited number (2) of donors in this study. While we observed many consistent responses between donors, we also observed some variability in the secretome of macrophage from discrete donors, and effect which has not been broadly characterized in literature. As a result we chose to not try to make definitive findings in regards to donor variability in this study. However, it will be important for our biomaterial community to pursue future projects to consider the effect of multiple donors to better understand the degree to which donor variability influence the temporal nature of macrophage polarization.

## 5. Conclusions

Craniofacial bone regeneration is a complex process involving the coordinated action of a heterogeneous mixture of cells, including progenitors and immune cells. The field of bone tissue engineering has largely concentrated on biomaterial designs to induce progenitor differentiation towards osteogenic phenotypes. As a result there is an opportunity to consider biomaterial designs that orchestrate patterns of pro-regenerative immune cell phenotypic trajectories. Here we describe the influence of pore architecture and glycosaminoglycan composition of a biomimetic mineralized collagen_GAG scaffold on the polarization of human macrophages towards a pro-healing phenotype. M0 macrophages could be effectively polarized towards pro-inflammatory (M1) and pro-healing (M2) phenotypes in mineralized collagen scaffolds via exogenous polarization factors in a manner similar to that seen in traditional 2D culture. Culture of M0 macrophages in mineralized collagen scaffolds absent any polarizing factors broadly induced an initial M1-like phenotype followed by progression towards an M2-like phenotype over the course of 7 days. Principal component analysis revealed this transition of M1-like to M2-like genomic phenotypes was temporally influenced by scaffold type, with surface antigen and secretome analyses providing additional insight about polarization kinetics. Macrophages in isotropic or anisotropic scaffolds with larger pore sizes showed greater M2-like surface antigen presentation with macrophages in anisotropic (aligned) scaffolds showing the greatest M2-like gene expression. Further, inclusion of chondroitin-6-sulfate (versus chondroitin-4-sulfate or heparin) increased M2-associated secretome and surface marker expression. This study suggests the potential for optimizing the structural and composition of mineralized collagen scaffolds to shape the inflammatory phenotype of scaffolds to further enhance regenerative potency.

## Acknowledgements

The authors would like to acknowledge the following institutes for access to their facilities and services: the School of Chemical Sciences Microanalysis Laboratory, the Carl R. Woese Institute for Genomic Biology, the Tumor Engineering and Phenotyping Shared Resource (TEP) at the Cancer Center at Illinois, and the Beckman Institute for Advanced Science and Technology, located at the University of Illinois. Research reported in this publication was supported by the National Institute of Dental and Craniofacial Research of the National Institutes of Health under Award Number R21 DE026582 and R01 DE030491 (BACH) as well as National Institute of Arthritis and Musculoskeletal and Skin Diseases under Award Number R01 AR077858 (BACH). We are also grateful for funds provided by the NSF Graduate Research Fellowship DGE-1746047 (VK) and the Chemistry-Biology Interface Research Training Program at the University of Illinois (T32 GM070421, VK). Additional support was provided by the Carl R. Woese Institute for Genomic Biology and the Chemical and Biomolecular Engineering Dept. at the University of Illinois at Urbana-Champaign. The interpretations and conclusions presented are those of the authors and are not necessarily endorsed by the National Institutes of Health or the National Science Foundation.

## Contributor Roles Taxonomy (CRediT) Contributions [51, 52]

**Vasiliki Kolliopoulos:** Conceptualization, Data curation, Formal Analysis, Visualization, Investigation, Methodology, Writing – original draft, Writing – review & editing. **Hashni E. Vidana Gamage:** Investigation, Formal Analysis. **Maxwell Polanek:** Investigation, Data curation, Formal Analysis, Writing – original draft, Writing – review & editing. **Melisande Wong Yan Ling:** Investigation. **Angela Lin:** Investigation. **Robert Guldberg:** Supervision, Methodology, Writing – review & editing. **Erik Nelson:** Supervision, Methodology, Writing – review & editing. **Kara Spiller:** Supervision, Methodology, Writing – review & editing. **Brendan Harley:** Conceptualization, Resources, Project administration, Funding acquisition, Supervision, Writing – review & editing.

## Disclosure

The authors have no conflicts of interest to report.

## Data availability

The raw data required to reproduce these findings are available upon request to Brendan Harley. The processed data required to reproduce these findings are available upon request to Brendan Harley.

## Notes

### Competing Interest Statement

The authors have declared no competing interest.

