## Supplemental Info for "Mineralized collagen scaffold pore architecture and glycosaminoglycan content biases anti-inflammatory macrophage phenotype"

[1] Dept. Chemical and Biomolecular Engineering,
University of Illinois at Urbana-Champaign, Urbana, IL 61801

[2] School of Molecular and Cellular Biology
University of Illinois at Urbana-Champaign, Urbana, IL 61801

[3] Phil and Penny Knight Campus for Accelerating Scientific Impact,
University of Oregon, Eugene, OR 97403

[4] Cancer Center at Illinois
University of Illinois at Urbana-Champaign, Urbana, IL 61801

[5] Dept. Biomedical Engineering, Science and Health Systems,
Drexel University, Philadelphia, PA 19104

[6] Carl R. Woese Institute for Genomic Biology
University of Illinois at Urbana-Champaign, Urbana, IL 61801

**Corresponding Author:**
B.A.C. Harley
Dept. of Chemical and Biomolecular Engineering
Cancer Center at Illinois
Carl R. Woese Institute for Genomic Biology
University of Illinois at Urbana-Champaign
110 Roger Adams Laboratory
600 S. Mathews Ave.
Urbana, IL 61801


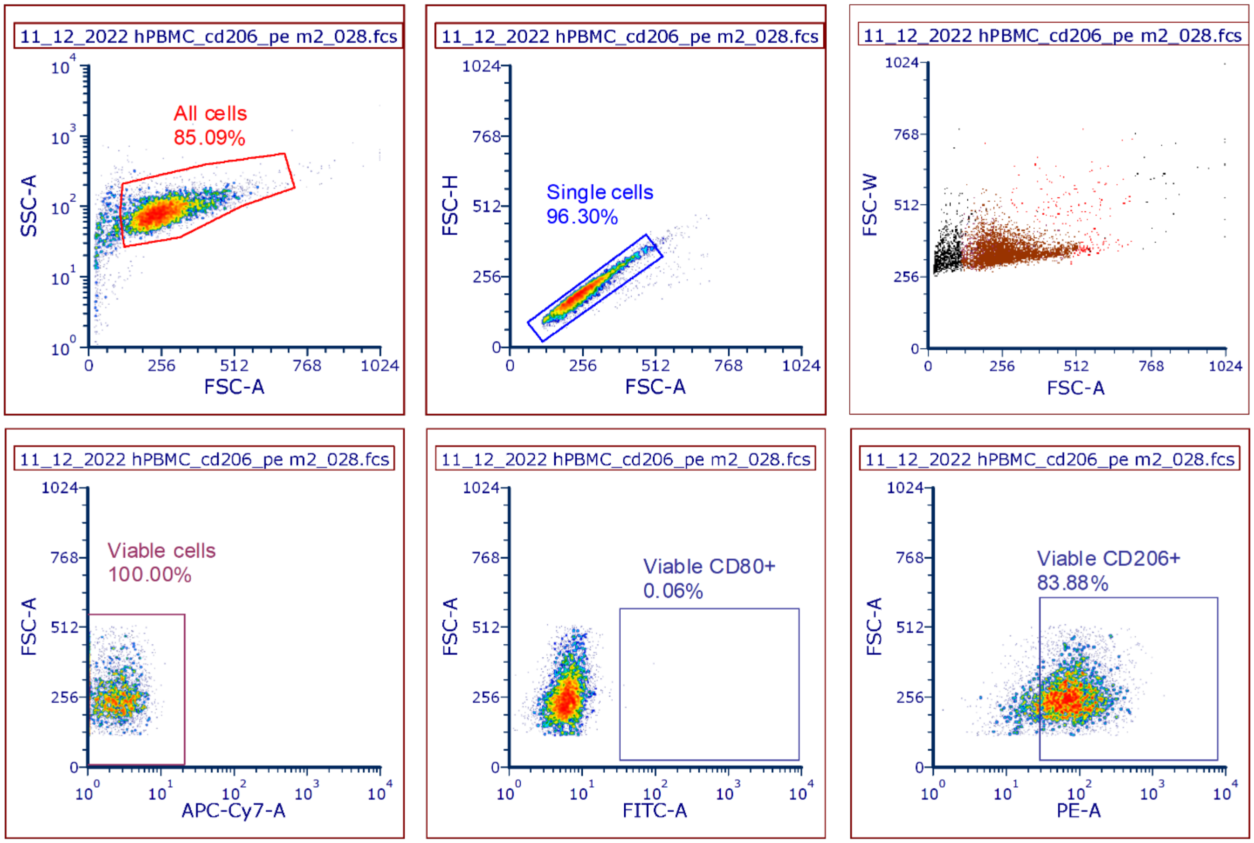


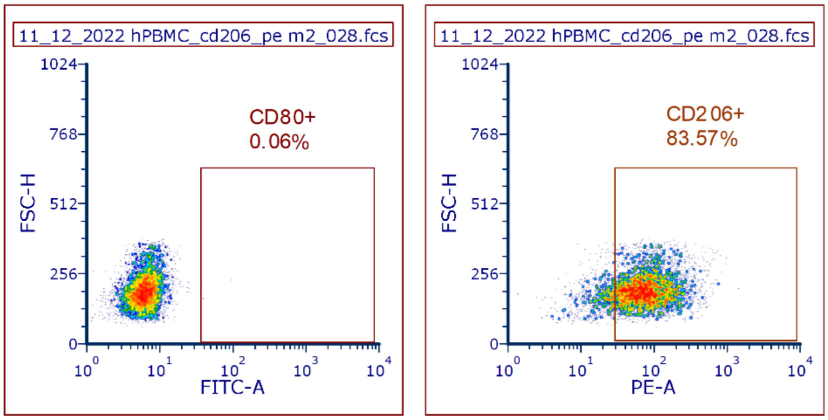


| Gate | # of Events | % of gated cells | % of gated - parent | % of all cells |
|---|---|---|---|---|
| None | 5204 | 100.00 | 96.30 | 81.94 |
| All cells | 5204 | 100.00 | 96.30 | 81.94 |
| Single cells | 5204 | 100.00 | 96.30 | 81.94 |
| CD80+ | 3 | 0.06 | 0.06 | 0.05 |
| CD206+ | 4349 | 83.57 | 80.48 | 68.48 |

**Supp. Figure 1:** Representative gating strategy for CD80+ and CD206+ cells.

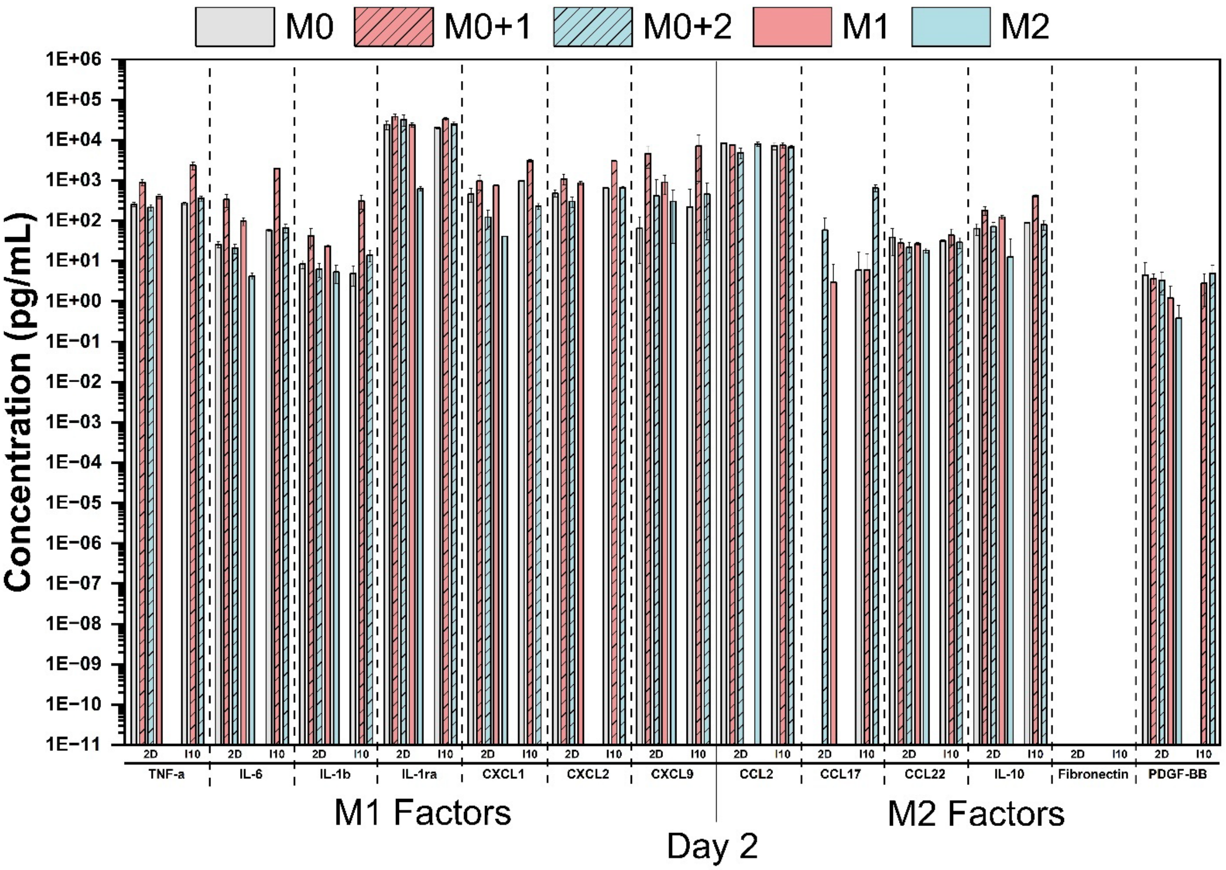


**Supp. Figure 2**: Custom Luminex panel of 7 M1 and 6 M2 soluble factors used to characterize the secretome of macrophages pre-polarized to M1 and M2 or cultured on 2D surfaces with and without polarizing cytokines as compared to 3D cultures. Samples were collected on day 2 of culture. Concentrations depicted on a log10 scale with units of pg/ml.

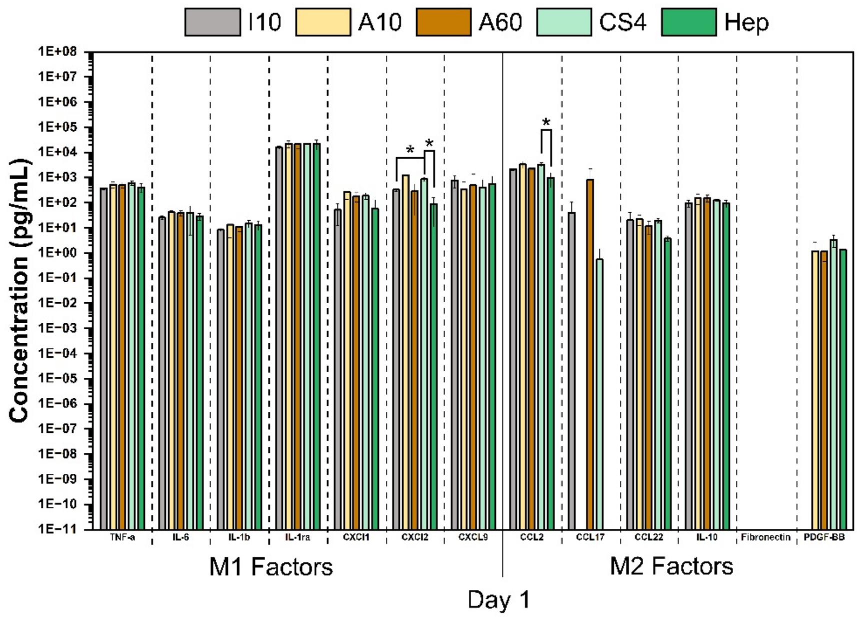


CXCL2: Hep had an outlier 0 value (not removed)

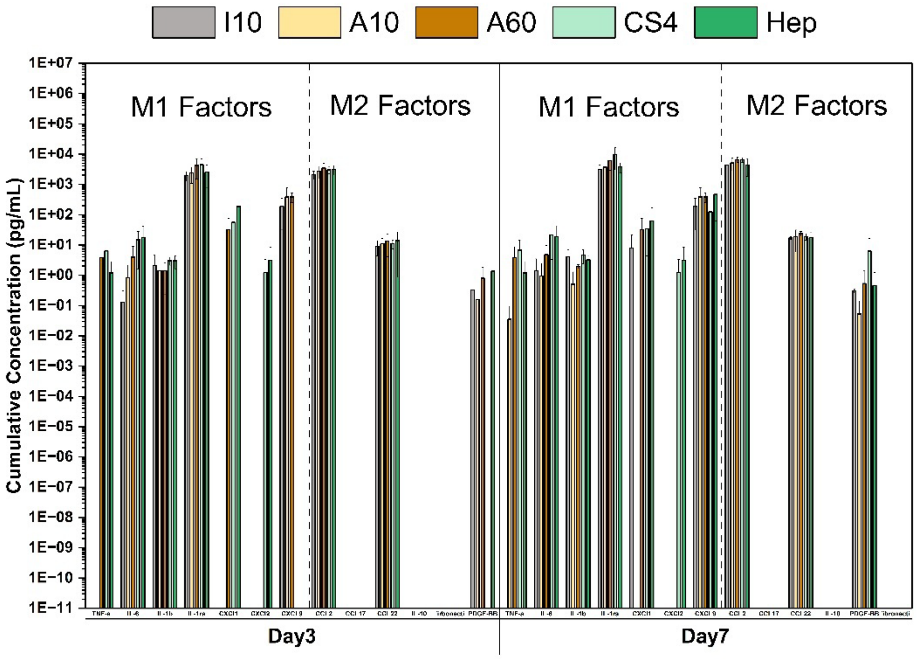


**Supp. Figure 3**: Custom Luminex panel of 7 M1 and 6 M2 soluble factors used to characterize the secretome of M0 macrophages cultured on mineralized collagen scaffolds with varying pore size and glycosaminoglycan content for 7 days. Concentrations (pg/ml) depicted on a log10 scale. Scaffold groups include isotropic -10 (I10), anisotropic -10 (A10), anisotropic -60 (A60) all with chrondroitin-6-sulfate, isotropic chondroitin-4-sulfate (CS4), and isotropic heparin (Hep). * indicates significance at p<0.05 between indicated groups at that timepoint and ^ indicates significance at p<0.05 of the indicated group and all other groups at that timepoint.

**Supp. Table 1. Detailed information about flow cytometry reagents used for macrophage characterization.** Fc blocker includes a panel of CD64, CD32, and CD16.

| Target | Species / Clone | Conjugate | Host / Isotype | Source | Dilution | Catalog # |
|---|---|---|---|---|---|---|
| CD80 (M1) | Human / CD10.4 | FITC | Mouse / IgG1κ | eBioscience | 100x | 11-0809-42 |
| CD206 (M2) | Human / 19.2 | PE | Mouse / IgG1κ | eBioscience | 100x | 12-2069-42 |
| Live/DEAD | N/A / N//A | Violet | N/A | Thermo Fisher | 1000x | L34955 |
| Fc Receptor Blocker | Human / N/A | N/A | Human | eBioscience | | 14-9161-71 |

**Supp. Table 2**: Custom Luminex panel factors

| **Macrophage phenotype** | **Long name** | **Short name** |
|---|---|---|
| M1 | Tumor necrosis factor alpha | TNF-α |
| | Interleukin 6 | IL-6 |
| | Interleukin 1b | IL-1b |
| | Interleukin 1ra | IL1-ra |
| | Chemokine ligand 1 | CXCL1 |
| | Chemokine ligand 2 | CXCL2 |
| | Chemokine ligand 9 | CXCL9 |
| M2 | Chemokine ligand 2 | CCL2 |
| | Chemokine ligand 17 | CCL17 |
| | Chemokine ligand 22 | CCL22 |
| | Interleukin 10 | IL-10 |
| | Fibronectin | Fibronectin |
| | Platelet derived growth factor subunit B | PDGF-BB |

**Supp. Table 3**: Genes examined via a custom Nanostring panel

| Gene name | Probe code | Function |
|---|---|---|
| ACTA2 | NM_001613.1 | ECM |
| AGGF1 | NM_018046.3 | Angiogenesis |
| ANG | NM_001145.4 | M2 |
| BGN | NM_001711.3 | ECM |
| BTG1 | NM_001731.2 | Immune Signaling |
| CABLES1 | NM_001100619.2 | M2 |
| CCL15 | NM_032965.4 | Angiogenesis |
| CCL17 | NM_002987.2 | M2 |
| CCL18 | NM_002988.2 | M2 |
| CCL2 | NM_002982.3 | M1 |
| CCL22 | NM_002990.3 | M2 |
| CCL24 | NM_002991.2 | M2 |
| CCL26 | NM_006072.4 | M2 |
| CCL5 | NM_002985.2 | Angiogenesis |
| CCL8 | NM_005623.2 | M1 |
| CCR7 | NM_001838.2 | M1 |
| CD163 | NM_004244.4 | M2c |
| CD200R1 | NM_138806.3 | M2 |
| CD80 | NM_005191.3 | M1 |
| CLEC10A | NM_182906.2 | M2 |
| COL1A1 | NM_000088.3 | ECM |
| COL3A1 | NM_000090.3 | ECM |
| COL5A1 | NM_000093.3 | ECM |
| CTGF | NM_001901.2 | ECM |
| CTNNB1 | NM_001098210.1 | Angiogenesis |
| CXCL12 | NM_199168.3 | M1 |
| CXCR4 | NM_003467.2 | Immune Signaling |
| DACT1 | NM_001079520.1 | M2 |
| DCN | NM_001920.3 | ECM |
| EGFL7 | NM_016215.3 | Angiogenesis |
| ETS1 | NM_005238.3 | Angiogenesis |
| FGF2 | NM_002006.4 | ECM |
| FLT1 | NM_002019.4 | Angiogenesis |
| FN1 | NM_212482.1 | ECM |
| FOXO1 | NM_002015.3 | Angiogenesis |
| FOXO3 | NM_001455.2 | Angiogenesis |
| FOXO4 | NM_001170931.1 | Angiogenesis |
| FST | NM_006350.2 | Angiogenesis |

| | | |
|---|---|---|
| FYN | NM_002037.3 | Immune Signaling |
| HSPG2 | NM_005529.5 | ECM |
| IDO1 | NM_002164.5 | M1 |
| IGF1 | NM_000618.3 | M2 |
| IL1B | NM_000576.2 | M1 |
| IL6 | NM_000600.3 | M1 |
| JAG1 | NM_000214.2 | Immune Signaling |
| LUM | NM_002345.3 | ECM |
| MARCO | NM_006770.3 | M2c |
| MMP2 | NM_004530.2 | M1 |
| MMP9 | NM_004994.2 | M2c |
| MRC1 | NM_002438.2 | M2 |
| PDGFA | NM_002607.5 | Angiogenesis |
| PDGFB | NM_033016.2 | Angiogenesis |
| PDGFC | NM_016205.2 | Angiogenesis |
| PDGFRA | NM_006206.3 | ECM |
| PDGFRB | NM_002609.3 | ECM |
| RAMP1 | NM_005855.2 | M2 |
| SPP1 | NM_000582.2 | Immune Signaling |
| STAT3 | NM_003150.3 | Immune Signaling |
| STAT6 | NM_003153.3 | Immune Signaling |
| TGFB1 | NM_000660.3 | ECM |
| TIE1 | NM_005424.2 | Angiogenesis |
| TIMP3 | NM_000362.4 | Immune Signaling |
| TNF | NM_000594.2 | M1 |
| TNFRSF11A | NM_003839.3 | M2 |
| VCAN | NM_004385.3 | ECM |
| VEGFA | NM_001025366.1 | Angiogenesis |
| VEGFB | NM_003377.3 | Angiogenesis |
| VEGFC | NM_005429.2 | Angiogenesis |
| VIM | NM_003380.2 | ECM |
| WNT5A | NM_003392.3 | Immune Signaling |
| GAPDH | NM_001256799.1 | Housekeeping |
| TBP | NM_001172085.1 | Housekeeping |
